# Comparative performance of four nucleic acid amplification tests for SARS-CoV-2 virus

**DOI:** 10.1101/2020.03.26.010975

**Authors:** Yujuan Xiong, Zhen-Zhen Li, Qi-Zhen Zhuang, Yan Chao, Fei Li, Yi-Yuan Ge, Yi Wang, Pei-Feng Ke, Xian-Zhang Huang

## Abstract

Coronavirus disease 2019 (COVID-19) can be screened and diagnosed through the detection of severe acute respiratory syndrome coronavirus 2 (SARS-CoV-2) by real-time reverse transcription polymerase chain reaction. SARS-CoV-2 nucleic acid amplification tests (NAATs) have been rapidly developed and quickly applied to clinical testing during the pandemic. However, studies evaluating the performance of these NAAT assays are limited. We evaluated the performance of four NAATs, which were marked by the Conformité Européenne and widely used in China during the pandemic. Results showed that the analytical sensitivity of the four assays was significantly lower than that claimed by the NAAT manufacturers. The limit of detection (LOD) of Daan, Sansure, and Hybribio NAATs was 3000 copies/mL, whereas the LOD of Bioperfectus NAATs was 4000 copies/mL. The results of the consistency test using 46 samples showed that Daan, Sansure, and Hybribio NAATs could detect the samples with a specificity of 100% (30/30) and a sensitivity of 100% (16 /16), whereas Bioperfectus NAAT detected the samples with a specificity of 100% (30/30) and a sensitivity 81.25% (13/16). The sensitivity of Bioperfectus NAAT was lower than that of the three other NAATs; this finding was consistent with the result that Bioperfectus NAAT had a higher LOD than the three other kinds of NAATs. The four above mentioned reagents presented high specificity; however, for the detection of the samples with low virus concentration, Bioperfectus reagent had the risk of missing detection. Therefore, the LOD should be considered in the selection of SARS-CoV-2 NAATs.

## Introduction

Severe acute respiratory syndrome coronavirus 2 (SARS-CoV-2) has spread rapidly since its recent identification in patients with severe pneumonia in Wuhan, China(1–3). To date, SARS-CoV-2 has affected more than 372,000 patients worldwide and resulted in more than 16,000 deaths (as of March 24, 2020)(4). The World Health Organization (WHO) has declared the Chinese outbreak of coronavirus disease 2019 (COVID-19) to be a Public Health Emergency of International Concern and stated that the spread of COVID-19 may be interrupted by early detection, isolation, prompt treatment, and the implementation of a robust system(4, 5). On March 11, at a regular press conference in Geneva, WHO Director-General Tan Desai said that the COVID-19 epidemic can be called a pandemic in terms of its characteristics. Unfortunately, no drug or vaccine has yet been approved to treat human coronaviruses, and new interventions are likely to require months to years to develop(6). Therefore, early diagnosis of SARS-CoV-2 infection is important to distinguish it from asymptomatic, healthy, and other pathogenic infections and to ensure timely isolation of the infected patients.

At present, various detection methods for SARS-CoV-2 infection have been reported, such as real-time reverse transcription–polymerase chain reaction (RT-PCR) assay, sequencing, CRISPR technique, nucleic acid mass spectrometry, and serum immunology(7). However, real-time RT-PCR remains the primary means for diagnosing COVID-19 among the various diagnostic platforms available. The development of RT-PCR methods to diagnose COVID-19 mainly targets various combinations of the open reading frame (ORF), envelope (E), nucleocapsid (N), spike (S), and RNA-dependent RNA polymerase (RdRp) genes(8–10).

Although real-time RT-PCR assay is a highly sensitive technique, false negative results have still been reported in some COVID-19 patients. These results may occur due to insufficient organisms in the specimen resulting from improper collection, transportation, storage, and handling, as well as laboratory test conditions and personnel operation(11–13). Moreover, the quality of the examined regents is related closely to the PCR results for SARS-CoV-2. In response to this outbreak, a number of nucleic acid amplification tests (NAATs) for SARS-CoV-2 were rapidly developed in China and quickly applied to clinical testing. To date, no quality evaluation of SARS-CoV-2 NAATs has been reported.

The purpose of this study was to comprehensively evaluate the performance of four SARS-CoV-2 NAATs, which were Conformité Européenne (CE) marked and widely used. The results of this study will be helpful for laboratorians to select the appropriate assay for detecting SARS-CoV-2.

## Materials and methods

### Tested samples

The kit of performance verification reference materials, which were cell cultures containing pseudoviruses, included 10 positive reference materials, two negative reference materials, 18 analytical specificity reference materials, and three interference reference materials (Guangzhou BDS Biological Technology Company Limited).The positive quality control agents were made with cell cultures containing SARS-CoV-2 pseudoviruses (Guangzhou BDS Biological Technology Company Limited).The pseudoviruses in the positive quality control agents included segments of the ORF1ab gene (the genome coordinates: 900–1500,12200–13500, 18770–18950, and 20560–24453), N gene, and E gene, which were the important characteristic genes of SARS-CoV-2. RNA transcripts, presented by the Chinese Academy of Metrology, were used as standard materials. The RNA transcripts obtained the complete N gene, the complete E gene, and ORF1ab gene segment (14911-15910, GenBank No. NC_O45512) of SARS-CoV-2. The standard value of reference material was obtained by absolute quantitative digital PCR. RNA was extracted from cell cultures with the nucleic acid extraction reagent (Tianlong Technology Company Limited), aliquoted into multiple tubes for testing with each NAAT method, and stored at −80 °C.

Human samples (n = 46), which were RNA leftovers from daily laboratory activity, were provided by the Guangdong Provincial Hospital of Chinese Medicine and Hybribio Medical Laboratory. The selection of clinical samples used in the study was exclusively based on their SARS-CoV-2 virus detection result. The study was approved by the Ethics Committees from the Guangdong Provincial Hospital of Chinese Medicine (ZE2020-027-01).

### Assay and equipment

Four sets of systems were used to detect the SARS-CoV-2 virus: Daan SARS-CoV-2 NAAT on LightCycler 480 II PCR instrument (Daan NAAT), Sansure SARS-CoV-2 NAAT on SLAN-96P PCR instrument (Sansure NAAT), Hybribio SARS-CoV-2 virus NAAT on SLAN-96P PCR instrument (Hybribio NAAT), and Bioperfectus SARS-CoV-2 NAAT on SLAN-96P PCR instrument (Bioperfectus NAAT). Real-time PCR was applied to the four systems, which targeted the combinations of the ORF1ab gene, N gene of SARS-CoV-2, and endogenous gene of humans. About 5 μL of RNA was added to the PCR mixture and tested according to the manufacturers’ instructions for each assay. Positive results were defined as the simultaneous detection of the ORF1ab gene and N gene. The four systems had different detection sites for the ORF1ab gene. The PCR primers designed for the four systems were complementary to the sites on the SARS-CoV-2 genomic sequence (GenBank No. NC_045512) in the region of 20700–21000 (Daan), 12500–13300 (Sansure), 13400–13550 (Bioperfectus), and 7100–7600 bp (Hybribio). Cycle threshold (Ct) values (i.e., number of cycles required for the fluorescent signal to cross the threshold in RT-PCR) quantified viral load, with low values indicating a high viral load.

### Analytical Sensitivity

Analytical sensitivity was examined in accordance with a modified EP17-A protocol, MM03 and MM19-A guidelines of Clinical Laboratory Standards Institute (CLSI). The analytical sensitivity of each NAAT assay was compared using the same RNA samples obtained from the extraction of mixed clinical samples or positive quality controls. The concentrations of RNA samples were determined by fluorescence quantitative method using standard materials as standard. Serial dilutions were made with a known concentration of the target substance in the analytical range of the expected limit of detection (LOD) and tested in replicates of 20. All panel members were prepared at the same time, aliquoted into individual tubes for each concentration and each assay, stored frozen, and thawed on the day of testing. When 19 or all 20 replicates were positive, the concentration was temporarily defined as LOD, followed by 20 replicates per day for two consecutive days, yielding a total of 60 results. If more than 95% of the results were positive, the concentration was finally determined as the LOD of the reagent. The experiments were spread over 3 days so that the standard deviations reflect the performance of the assay over a range of typical laboratory conditions but without a change in reagent lots.

### Precision

Precision was evaluated in accordance with a modified EP12 protocol, MM09 and MM19-A guideline of CLSI using a patient RNA pool with concentration above 20% of the LOD and the negative sample. Sample pools were divided into five aliquots per level and frozen at −70 °C. One aliquot of each level was thawed daily and analyzed four times per day during a period of five consecutive workdays (n = 20 per level). Precision was evaluated as the coefficient of variation (CV), which was calculated from the data series mean and standard deviation.

### Accuracy

Ten positive reference materials and two negative reference materials from the performance verification reference material kit were used to evaluate accuracy.

### Analytical specificity

Eighteen pseudovirus samples of analytical specificity reference materials from the performance verification reference material kit were used to determine the analytical specificity of four NAATs, including human coronavirus HCoV-OC43, HCoV-HKU1 RNA, HCoV-229E RNA, HCoV-NL63 RNA, SARS RNA, Middle East respiratory syndrome (MERS) RNA, influenza A HIN1 virus, influenza B INFB virus, respiratory syncytial virus type A and type B, human parainfluenza virus, adenovirus, enterovirus, mycoplasma pneumoniae, Epstein–Barr (EB) virus, human cytomegalovirus, *Mycobacterium tuberculosis,* and two samples with human genome DNA.

### Analytical interferences

To investigate the analytical interferences, three interference reference materials from the performance verification reference material kit were used, including 6 g/dL hemoglobin, 30 g/dL albumin, and the mix of 100 μg/mL ribavirin and 100 μg/mL azithromycin. All tests were repeated three times.

### Method comparison

Forty-six RNA samples from suspected patients with COVID-19 were used in the method comparison. Given that no reference method was used to determine the sensitivity and specificity of NAATs, the consensus result was determined for each sample; samples were categorized as true positive (TP) if positive by two or more assays (14). The SARS-CoV-2-positive samples were further confirmed by sequencing.

### Statistical analysis

The significance of the difference in sensitivity between the NAATs was assessed by using Fisher’s exact test in GraphPad Prism (5.0). The Chi-square likelihood ratio test was used to analyze the discordant results. The 95% confidence intervals (CIs) for specificity and sensitivity were calculated by using the Wilson–Score method in GraphPad Prism. Significant differences were considered at *P* values less than 0.05.

## Result

### Analytical sensitivity

We evaluated the analytical sensitivity of each assay to determine differences in the ability of each assay to detect SARS-CoV-2 in mixed clinical samples or in pseudovirus cultures. First, we contrived RNA specimens from SARS-CoV-2 pseudovirus cultures at four separate final concentrations (500, 1000, 2000, and 3000 copies/mL). The ORF1ab gene test results of the serial dilutions were all negative by the Hybribio and the Bioperfectus NAATs. Further investigation revealed that the pseudoviral RNA did not contain the detecting locus of Hybribio and Bioperfectus NAATs targeting the ORF1ab gene. Daan NAAT detected 18/20 replicates at 2000 copies/mL and 20/20 replicates at 3000 copies/mL. Sansure NAAT detected 17/20 at 1000 copies/mL and 20/20 at 2000 copies/mL (Table 1). The final results of 60 replicates showed that the LOD of Daan NAAT was 3000 copies/mL for pseudovirus cultures, and the LOD of Sansure NAAT was 2000 copies/mL for pseudovirus cultures.

**Table 1.** Comparison of Analytical sensitivity.

| Copies/mL | Pseudoviruses |  |  |  | Clinical samples |  |  |  |
| --- | --- | --- | --- | --- | --- | --- | --- | --- |
|  | Daan | Sansure | Hybribio | Bioperfectus | Daan | Sansure | Hybribio | Bioperfectus |
| **4000** | / | / | / | / | 20/20 | 20/20 | 20/20 | 20/20 |
| **3000** | 20/20 | 20/20 | 0/20 | 0/20 | 20/20 | 20/20 | 20/20 | 12/20 |
| **2000** | 18/20 | 20/20 | 0/20 | 0/20 | 12/20 | 17/20 | 17/20 | 5/20 |
| **1000** | 12/20 | 17/20 | 0/20 | 0/20 | / | / | / | / |
| **500** | 5/20 | 17/20 | 0/20 | 0/20 | / | / | / | / |

To explore the LOD of the NAATs in clinical samples, we further contrived mixed RNA specimens from clinical samples at three separate final concentrations (2000, 3000, and 4000 copies/mL). Daan NAAT detected 12/20 replicates at 2000 copies/mL and 20/20 replicates at 3000 copies/mL (Table 1). Sansure NAAT detected 17/20 at 2000 copies/mL and 20/20 at 3000 copies/mL. Hybribio NAAT detected 10/20 replicates at 2000 copies/mL and 20/20 replicates at 3000 copies/mL. Bioperfectus NAAT detected 10/20 at 3000 copies/mL and 20/20 at 4000 copies/mL. The final results of 60 replicates showed that the LOD of clinical samples was 3000 copies/mL by Daan, Sansure, and Hybribio NAATs, whereas that of clinical samples was 4000 copies/mL by Bioperfectus NAAT. The LODs declared in the instructions of Daan, Sansure, Hybribio, and Bioperfectus NAATs were 500, 200,1000, and 1000 copies/mL, respectively.

### Precision

For the negative sample, the 20 repeated test results by all four NAATs had no amplification curve of target genes but a positive amplification curve of the internal reference gene. For the positive sample, 20 repeated test results by all four NAATs had positive amplification curves of the target genes and internal reference gene. The within-run CVs at the concentrations above the 20% level of the LOD on all four systems were less than 4%, and the total CVs were less than 5% (Table 2).

**Table 2.**
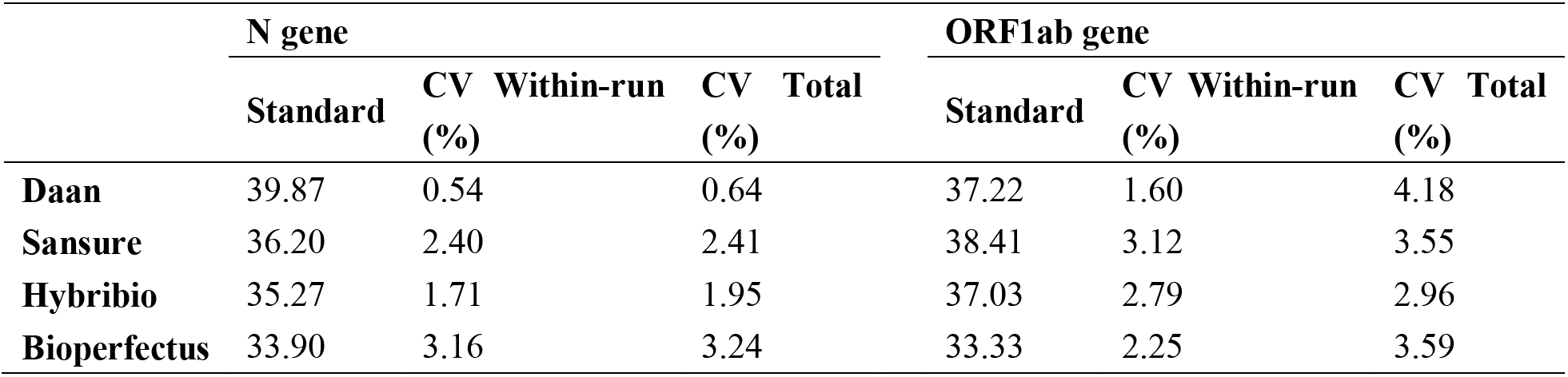
Comparison of precision.

### Accuracy

The results of all four NAATs for accuracy evaluation were in line with expectations by using 10 positive reference materials and two negative reference materials.

### Analytical specificity

For the four assays, no amplification curves were obtained for up to 10,000 copies/mL of six coronavirus pseudovirus samples with HCoV-OC43, HCoV-HKU1, HCoV-229E, HCoV-NL63, SARS, and MERS RNA. For the 10 common respiratory pathogens and two human genomic DNA samples, all results of the four assays were also negative.

### Analytical interferences

An assessment of interferences is shown in Table 3. The bias of Ct value for the N gene and ORF1ab gene on all four systems was less than 8.5% at the following interfering substance concentrations tested: 6 g/dL hemoglobin, 30 g/dL albumin, 100 μg/mL ribavirin, and 100 μg/mL azithromycin.

**Table 3.** Sample interferences for common interferents.

| Interferent | Relative bias from native sample, Ct% |  |  |  |  |  |  |  |
| --- | --- | --- | --- | --- | --- | --- | --- | --- |
|  | Daan |  | Sansure |  | Hybribio |  | Bioperfectus |  |
|  | N | ORF1ab | N | ORF1ab | N | ORF1ab | N | ORF1ab |
| **Hemoglobin** | 7.29 | 5.96 | 6.06 | 2.84 | 7.45 | 7.00 | 7.20 | 4.15 |
| **Albumin** | 6.72 | 5.92 | 6.83 | 3.39 | 8.22 | 7.96 | 7.20 | 3.72 |
| **Ribavirin & Azithromycin** | 6.72 | 5.92 | 6.76 | 4.54 | 7.86 | 7.60 | 7.20 | 4.23 |

### Comparison of the four NAATs

We defined consistency as two or more tests that produce same results and used these results to define true positives and negatives. By this definition, 34.8% (16/30) tested positive and 65.2% (30/46) tested negative. The results for all four NAATs were negative for all 30 negative samples, and the 16 positive samples were reported positive by Daan, Sansure, and Hybribio NAATs. However, three of the 16 positive samples were reported negative by Bioperfectus NAAT (Table 4). The Ct values of the N gene and ORF1ab gene in the three positive samples by Daan, Sansure, and Hybribio NAATs are shown in Table 5, and the Ct values of the ORF1ab gene were close to the positive threshold of the three NAATs declared in their instructions.

**Table 4.** Comparison of NAATs to consensus results for detection of SARS-CoV-2.

| Assays | TP | TN | FP | FN | Sensitivity (95% CI) | Specificity (95% CI) |
| --- | --- | --- | --- | --- | --- | --- |
| **Daan** | 16 | 30 | 0 | 0 | 100%(80.64 - 100) | 100%(80.64 - 100) |
| **Sansure** | 16 | 30 | 0 | 0 | 100%(80.64 - 100) | 100%(80.64 - 100) |
| **Hybribio** | 16 | 30 | 0 | 0 | 100%(80.64 - 100) | 100%(80.64 - 100) |
| **Bioperfectus** | 13 | 30 | 0 | 3 | 81.25%(56.99 - 93.41) | 100%(88.65 - 100) |

**Table 5.**
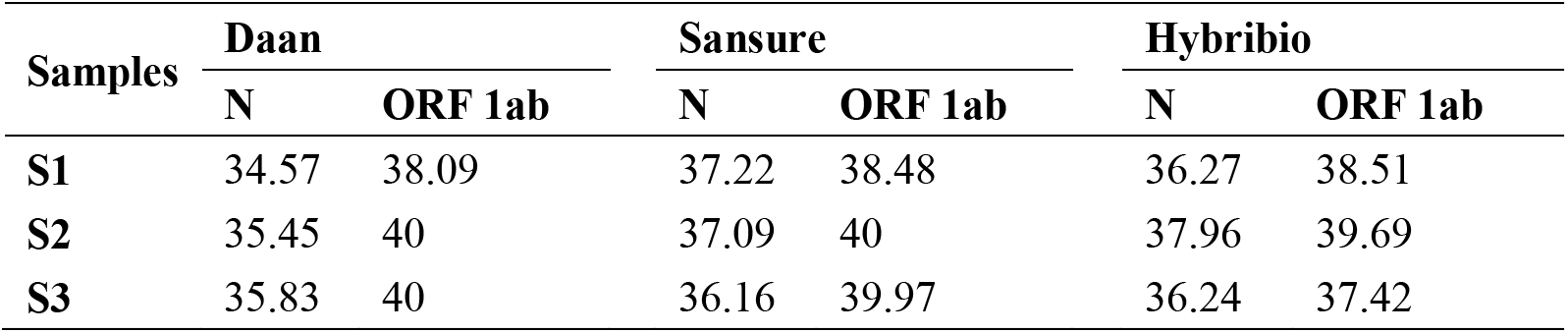
Ct values of the three clinical samples which was SARS-CoA-2 negative detected by Bioperfectus NAATs.

| Samples | Daan |  | Sansure |  | Hybribio |  |
| --- | --- | --- | --- | --- | --- | --- |
|  | N | ORF 1ab | N | ORF 1ab | N | ORF 1ab |
| **S1** | 34.57 | 38.09 | 37.22 | 38.48 | 36.27 | 38.51 |
| **S2** | 35.45 | 40 | 37.09 | 40 | 37.96 | 39.69 |
| **S3** | 35.83 | 40 | 36.16 | 39.97 | 36.24 | 37.42 |

The sensitivity and specificity for each NAAT were calculated based on consistent analysis. For Daan, Sansure, and Hybribio NAATs, their specificity and sensitivity were 100%. For Bioperfectus NAAT, the sensitivity was 81.25% and specificity was 100%. No significant difference was found in the sensitivity between Bioperfectus NAAT and the three other NAATs (P > 0.05).

## Discussion

The prevention of the COVID-19 epidemic is very grim. On the basis of China’s experience in preventing the further spread of the disease in the past 2 months, identifying COVID-19 from patients with other diseases as soon as possible has been proven to be particularly important to execute early isolation and early treatment (4, 5, 15). The use of RT-PCR technology to detect SARS-CoV-2 nucleic acid is an effective means to screen SARS-CoV-2 virus-infected patients. The current SARS-CoV-2 NAATs, which are widely used in China, were developed in response to the emergency situation of screening for SARS-CoV-2 infection. The performance of the assays is exactly what all testers want to know. In this study, four NAATs were evaluated. The results showed that the LODs of the four assays were significantly higher than the LODs declared in their instructions, suggesting that each laboratory should re-evaluate the LOD of the reagents depending on the needs of their laboratory. The RNA detection data from pseudovirus cultures showed that the LOD of Sansure was slightly lower than that of Daan NAAT. The RNA detection data from clinical samples showed that the LOD of Bioperfectus NAAT was slightly higher than that of the three other reagents but still reached the detection sensitivity of 20 copies per PCR reaction. There was no significant difference in the LOD of SARS-CoV-2 virus between pseudovirus cultures and clinical samples for Daan and Sansure NAATs. Thus, pseudovirus cultures could be used as an alternative for clinical samples in the performance evaluation of LOD. In addition, the four assays showed high precision, and the CV value was less than 5% when the concentration was 20% higher than the LOD. There were no cross-reactions by the four assays with four common human coronaviruses, SARS, and MERS viruses and no cross-reaction with other common respiratory pathogens, such as respiratory syncytial virus, adenovirus, influenza virus, and *Mycoplasma pneumoniae.*

In the present study, we further compared four NAATs for the detection of SARS-CoV-2 in clinical samples from suspected COVID-19 patients. Results showed that the clinical sensitivity and specificity of Daan, Sansure, and Hybribio NAATs were 100%, while three of the positive samples were missed by Bioperfectus NAAT. The Ct values of the ORF1ab gene of the three samples tested with Daan, Sansure, and Hybribio reagents were all close to 40, indicating that the concentration of the three samples may be lower than the LOD of Bioperfectus. Therefore, the LOD should be included in the reference index when selecting reagents for SARS-CoV-2 in the laboratory. Sequencing verification of the PCR-positive samples in this study suggested that the four assays did not show false positives in the detection of limited clinical samples. Although the small clinical sample size limited the optimal evaluation of the performance of these assays, the results of this study can still help laboratories to select an appropriate SARS-CoV-2 NAAT supplier and interpret their detection results.

Obtaining high-quality viral RNA from original clinical samples is crucial to ensure the accuracy of the detection results by NAATs. Therefore, the poor quality of extraction reagents and the unoptimized extraction process of viral RNA would affect the detection results. In this study, the performance of NAATs was evaluated by using the same viral RNA samples. Therefore, the difference in the experimental results only reflected the difference in the performance of nucleic acid amplification for NAATs but not the diversity in the performance of the whole SARS-CoV-2 detection system, which usually includes the nucleic acid extraction system.

In addition, RNA viruses may show substantial genetic variability. This could result in mismatch between the primer and probes with the target sequence, which can diminish the assay performance or result in false negative results. Therefore, the manufacturer of NAATs should pay attention to the sequence update information of the virus database at any time to ensure no mutation exists in the genome sequences corresponding to the primers, especially for the sequences corresponding to the 3’ end of the primers.

In summary, the four SARS-CoV-2 NAATs commonly used with commercial assays showed good reliability, but the actual LOD was significantly higher than the declared LOD. The analytic sensitivity of Daan, Sansure, and Hybribio NAATs was slightly higher than that of Bioperfectus NAAT.

## Acknowledgments

The authors would like to thank the Chinese Academy of Metrology, Sansure, Bioperfectus, and Hybribio for supplying the test kits.

